# Neural substrates of navigational decision-making in *Drosophila* larva anemotaxis

**DOI:** 10.1101/244608

**Authors:** Tihana Jovanic, James W. Truman, Marc Gershow, Marta Zlatic

## Abstract

Small animals use sensory information to navigate their environment in order to reach more favorable conditions. in gradients of light, temperature, odors and CO_2_, *Drosophila* larvae alternate periods of runs and turns, regulating the frequency size and direction of turns, to move in a favorable direction. Whether larvae use the same strategies when navigating in response to somatosensory input is unknown. Further, while many of the sensory neurons that mediate navigation behaviors have been described, where and how these navigational strategies are implemented in the central nervous system and controlled by neuronal circuit elements is not well known. Here we characterize for the first time the navigational strategies of Drosophila larvae in gradients of air-current speeds using high-throughput behavioral assays and quantitative behavioral analysis. We find that larvae extend runs towards favorable directions and shorten runs in unfavorable directions, and that larvae regulate both the direction and amplitudes of turns. These results suggest similar central decision-making mechanisms underlie navigation behaviors in somatosensory and other sensory modalities. By silencing the activity of individual neurons and very sparse expression patterns (2 or 3 neuron types), we further identify the sensory neurons and circuit elements in the ventral nerve cord and brain of the larva required for navigational decisions during anemotaxis. The phenotypes of these central neurons are consistent with a mechanism where the increase of the turning rate in unfavorable conditions and decrease in turning rate in favorable conditions are independently controlled. In addition, we find phenotypes that suggest that the decisions of whether and which way to turn are controlled independently. Our study reveals that different neuronal modules in the nerve cord and brain mediate different aspects of navigational decision making. The neurons identified in our screen provide a basis for future detailed mechanistic understanding of the circuit principles of navigational decisionmaking.

## Introduction

Orientation behavior allows animals to move in the environment as a function of sensory information to find more favorable conditions. This behavior is essential for survival and is shared across the animal kingdom. Many small organisms navigate their environments and move towards more favorable conditions by biasing their motor decisions as a function of the changes in the sensory information [1-11].

Organisms like *C. elegans* and larval *Drosophila* were shown to alternates periods of forward movements with reorientation events during which they make directional changes (decisions). Typically, in *Drosophila* larvae, navigation involves two stereotyped motor patterns: runs, which are periods of forward crawling and turns which are reorientation events involving head sweeps (one or multiple) followed by a choice of direction [3-6, 9]. It is thought that larvae integrate sensory information during a run and decide to turn when the sensory environment become unfavorable, while integration of sensory information during a head sweep determines the direction of the turn [4, 6, 9]

Larvae use these same strategies when navigating gradients of odors (chemotaxis), CO_2_, light (phototaxis) and temperature (thermotaxis).

The sensory neurons and receptors that mediate navigation in some types of sensory modalities (odor, temperature, light, C0_2_) in Drosophila larvae have been extensively investigated [1, 5, 7-9, 12, 13, 14] and the computations and the behavioral dynamics of taxis behaviors in some sensory gradients were described in recent years using quantitative analysis methods [4-6, 8, 13-15]. However, the central components of the neural circuits underlying navigational decisions (when to turn, how much to turn and which way to turn) with the exception of recent studies that discovered types of neurons in the brain and the SEZ (suboesophagial zone) required for taxis behaviors [4, 16, 17], remain largely unknown. Also, whether these circuit elements are common to navigational decision-making in different sensory modalities is still a mystery.

Adult flies orient in response to currents of air during flight and in the context of response to odor plumes [18-20]. Orientation in response to a current of air is called anemotaxis (from the greek *anemo-άνεμο* for wind and taxis - 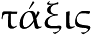 for arrangement, order). It is unknown whether *Drosophila* larvae can perform anemotaxis, and if so, if they do so via the strategies used by the chemosensory, thermosensory and photosensory circuits.

Here we investigate whether Drosophila larvae anemotax and show that larvae move away from high wind speeds. We characterize for the first time the navigational strategies of *Drosophila* larvae in a gradient of air-current speeds. We used a high-throughput behavioral assay combined with quantitative behavioral analysis to uncover the strategies the larva uses to navigate towards weaker wind speed areas. We show that the larva uses similar strategies to the ones it uses to navigate environments with varying concentrations of odors, gradients of CO_2_, light intensities and temperatures: they crawl forward in straight runs that are interrupted with reorientation turns. In the face of unfavorable changes in sensory input they increase their turn rate and the magnitude of those turns in order to more favorable conditions.

We then combined the high-throughput quantitative behavioral analysis methods with manipulation of neuronal activity (silencing using Tetanus toxin) and determined a sensory neuron type that mediates anemotaxis. In a targeted behavioral inactivation screen, we further identified 9 central neuron lines with very sparse expression patterns (1-3 neuron types) that drive in neurons involved in navigational decisionmaking during anemotaxis. We find that these neurons are located in the somatosensory circuitry (in the ventral nerve cord -VNC) and the brain and some of them implement specifically only certain types of navigations decisions, findings consistent with a modular organization of navigation-decision-making. These neurons represent the starting points for determining the circuit mechanisms underlying navigational decision-making.

## Results

### Anemotaxis -navigation in a gradient of air-current speeds

To determine how *Drosophila* larvae navigate in response to air currents, we presented fixed spatial gradients of wind speed to large numbers of animals while tracking their motion to analyze their behavioral dynamics.

We generated gradients of air-current speeds with one end of the arena at high wind speed and the other low. One gradient was between 3 m/s and 1 m/s and a second between 5 m/s and 2 m/s. We put larvae in the center of the arena and monitored their behavior for 10 minutes. We found that at the end of the experiment the majority of the larvae were located at or near the lower speed end, meaning that they go down the gradient of air-speed and away from stronger winds in an air-current speed gradient (Figure 1A).

**Figure 1.**
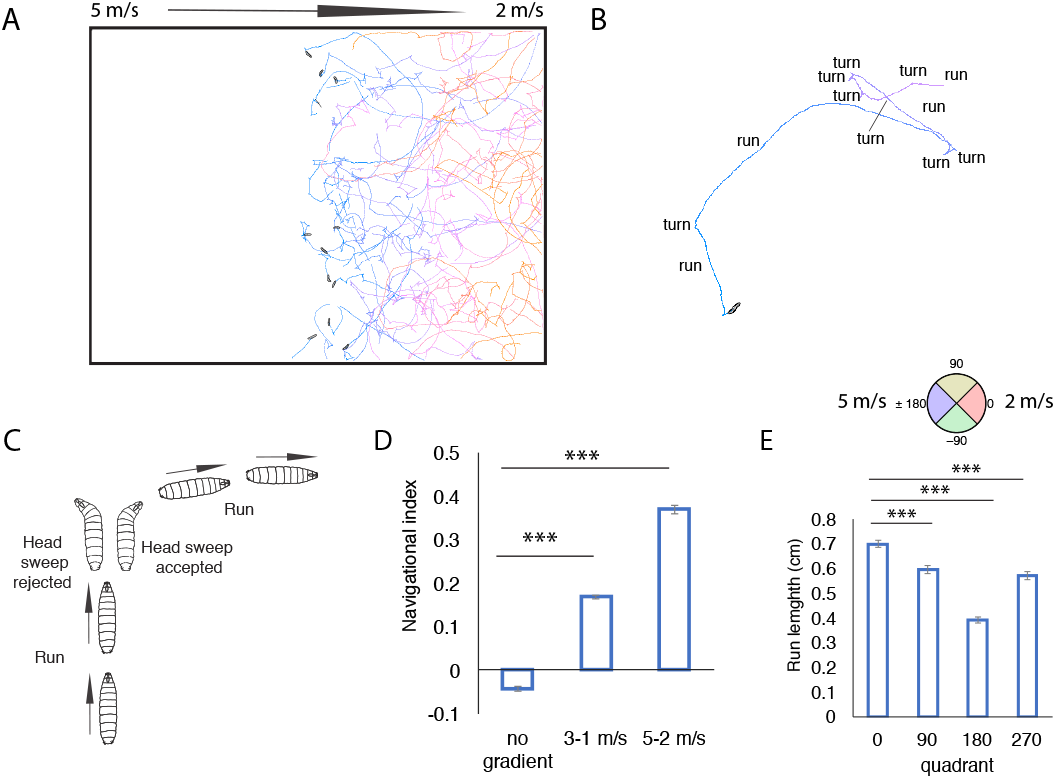
In air-speed gradient, the larvae navigate down the gradient. The colors of the tracks represent the time from the beginning of the experiment (blue) to the end (orange). Snapshots of the initial positions of larvae in the center of the agar plate are shown B Larvae alternate periods of runs and turns during which they sweep their heads and sample the sensory environment. C. An example of a reorientation event where a larva perpendicular to the direction of the gradient accepts a head sweep and extends a run in a favorable direction is shown D. Navigational index in 3-1 m/s and 5-2 m/s gradient. E. Run length in different quadrants (0, 90, 180, 270). The quadrtants are shown in the compass above the plot

During the assay, the larvae are put in the center of each agar plate in a single line along the y axis, perpendicular to the direction of the gradient of speed along the x axis. To quantify the overall navigational performance of Drosophila larvae in an air-speed gradient we computed the navigational index as a measure of the navigational response to the sensory gradient. The navigational index is computed by dividing the mean velocity of all larvae in the x direction, 〈*v_x_*〉, by the mean crawling speed, 〈*s*〉

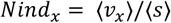

A navigational index of +1 would correspond to all larvae moving straight down the gradient, a navigational index of -1 to all the larvae moving straight up the gradient and a navigational index of 0 to larvae moving without bias towards 0 or 180 (down or up the gradient).

The navigational index was 0.17 in a 3 to 1 m/s gradient and 0.37 in a 5 to 2 m/s gradient. In a control experiment with no air-current at all, the navigation index was close to 0 as expected (Figure 1 D, Supplementary table 1). Larvae navigate away from the air-current and towards lower speeds in both the 3 to 1 m/s and 5 to 2 m/s

### Navigational strategies in anemotaxis

In order to uncover the behavioral strategies that the larvae use to navigate in gradients of air-speeds, we monitored 783 intact attP2>TNT larvae’s movements in the arena with a previously described quantitative behavioral analysis method [4, 5, 8]. This method quantifies the navigational performances of individual *Drosophila* larvae.

Navigational statistics across populations of larvae were further gathered to determine the behavioral strategies that the larvae use during taxis to bias their trajectories towards the areas of the arena with favorable conditions (in the case of anemotaxis, lower speeds of air-currents)

As for other types of taxis behaviors [3-6, 8, 9], during navigation larvae alternate periods of runs, which consists of forward crawling in approximately straight trajectories with periods of turns which allow them to change the direction of heading (Figure 1 A-C). We therefore, examined what biases in the runs interspersed with turns allowed the larvae to navigate the gradient towards the weaker wind speed areas of the arena (Figure 2, Supplementary figure 1).

**Figure 2.**
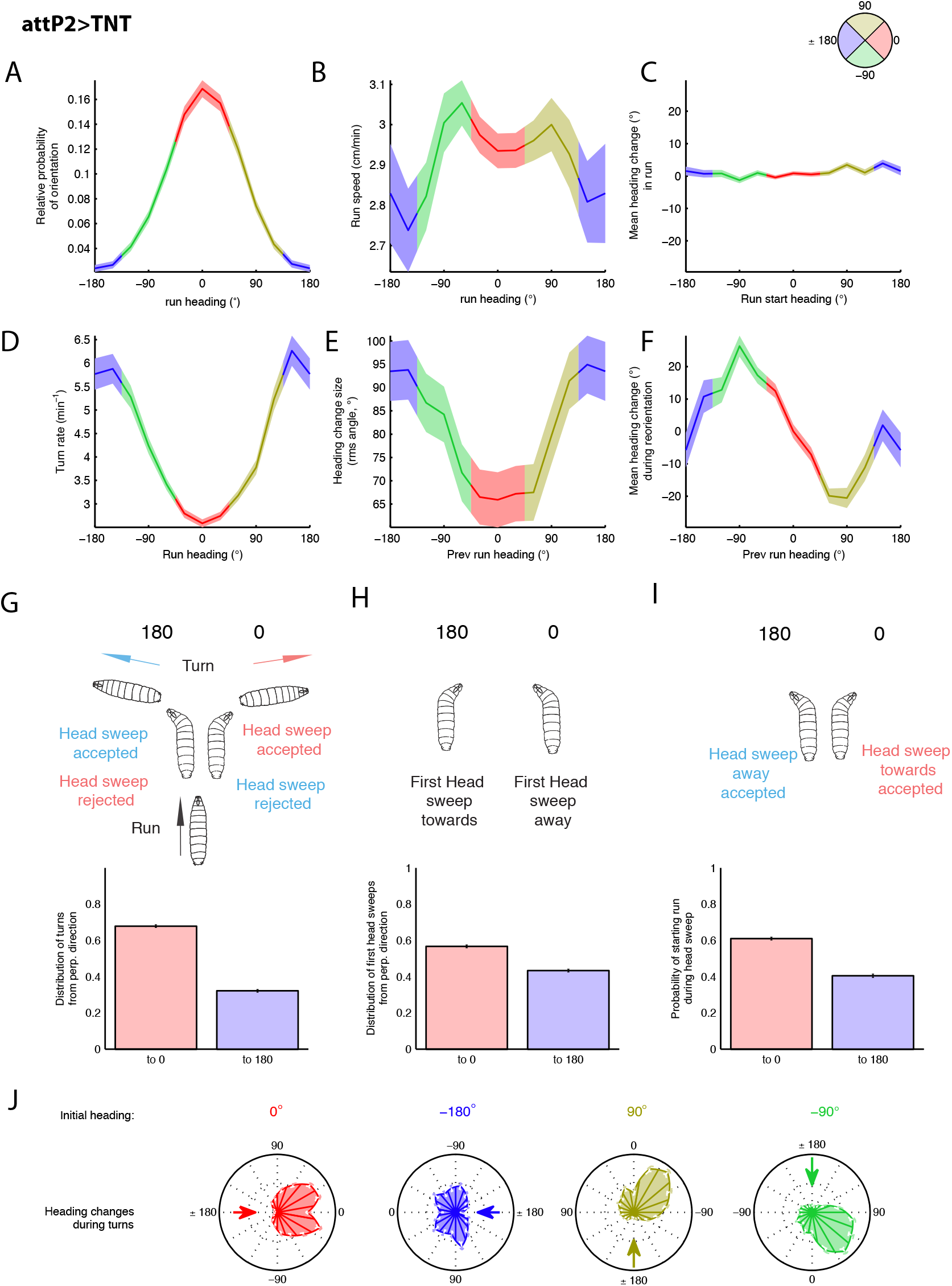
Navigational strategies in anemotaxis in control attP2>TNT (the same is shown for attP2-attP40>TNT in Supplementary figure 1) A. Relative probability of headings during runs. B. Speed versus heading during runs C. Mean heading change in runs D. Turn rate versus heading E. Turn size versus heading F. Mean heading change during reorientation G. Distribution of turns from perpendicular direction H. Distribution of head sweeps from perpendicular direction I. Probability of starting a run during a headsweep J. Heading changes during turns sorted by initial heading. Values are mean and s.e.m

A compass in which 0 indicates the direction down the gradient and 180 up the gradient was used to keep track of larval direction during runs and turns as a function of the wind speed spatial gradient.

It has been previously shown for other types of taxis, that larvae extend runs in favorable directions. We therefore measured how the length of runs depended on their directions, using 4 quadrants as shown in Figure 1C. We find that the run length is significantly longer in the 0 quadrant compared to the other three quadrant in the arena (Figure 1 E)

The crawling of larvae during runs is described by the magnitude (run speed) and direction (run heading) of the velocity vector. We calculated the fraction of time that larvae spent crawling in different directions and found that larvae spent the most time moving down air-speed gradients. (Figure 2 A). Larvae crawled slightly slower when heading up air-speed gradients than down (Figure 2 B).

We examined the rate at which larvae turned as a function of heading on a linear spatial gradient and found that the turn rate was highest when the larvae were heading towards high air-current speeds (180) and lowest when heading in the direction of the low speed end (Figure 2 D). As described for other types of taxis, larvae decrease their turn rate when they are headed in a favorable direction and increase their turn rate when headed in an unfavorable direction.

To examine whether larvae modulate the amplitude of turns to increase the probability of runs in a favorable direction, we examined the heading change effected by each turn, as a function of heading prior to the turn (Figure 2E). We found that larvae tend to make larger turns (average 93degree change in direction) when previously headed towards the wind source and smaller turns (average 65 degrees) when headed downwind. We next considered the average change in direction effected by each turn, as a function of prior heading (Figure 2F). We found that larvae biased the direction of their turns to move towards the downwind direction. In contrast, we found no bias in heading changes during runs (Figure 2C). We next examined the distribution of turn angles (Figure 2J). When larvae turned after a run up or down air-speed gradients the heading change distributions were bimodal and roughly symmetric for both direction, but they were narrower when larvae were initially headed in the favorable direction, consistent with smaller heading changes from the favorable direction (Figure 2J). When larvae turned after a run oriented perpendicular to the gradient, they did so with the same distribution of angular sizes to the left or right, but made more turns toward the favorable direction. This is consistent with a high probability of turning towards favorable directions. Indeed, when we quantified the probability of turns towards lowers speed end (0) versus high speed ends (180) of all the larvae after orthogonal to air-speed gradients, we found that nearly 68% of turns are towards the lower speed end (Figure 2 G).

During a turn, a larva sweeps its head to one side, after which it either starts a new run or initiates a new head sweep. To uncover bias in these head sweeping movements, we analyzed the statistics of all head sweeps initiated by larvae after runs pointed orthogonal to air-speed gradients. We found that the direction of the initial head sweep in each turn was biased towards the lower air-speed end (Figure 2H). In gradients of odors or CO2 larvae don’t show a significant bias of the first head sweep [5].

In those studies, second instar larvae were used, while here we use third instar *Drosophila* larvae. In a study that used third instar larvae in a chemotaxis assay, a bias of the first head sweep towards higher odor concentration was observed [6].

We next examined the biases in accepting a head sweep depending on its direction (whether the head sweep is towards the low speed end (0) or the high-speed end (180) and found that there is a significantly higher probability of accepting a head sweep when facing the lower speed end from the perpendicular direction (p<0.0001) (Figure 2 I)

In summary, during anemotaxis larvae use similar strategies as those previously described for other types of sensory gradient navigation: they modulate their turn rate, amplitude and direction so that they extend runs more when facing the favorable direction [4-6, 14]

### Chordotonal sensory neurons mediate anemotaxis

We have previously identified the chordotonal sensory neurons on the body wall of the larvae as key sensory neurons for sensing air-current [21, 22] [23] in a behavioral assay with uniform speeds in the arena. Here, we asked whether these sensory neurons also mediate anemotaxis of Drosophila larvae. We found that when silencing the chordotonal sensory neurons (air-current sensing neurons) the larvae perform taxis less efficiently as they navigational index was significantly reduced compared to attP2>TNT larvae (Figure 3 A).

**Figure 3.**
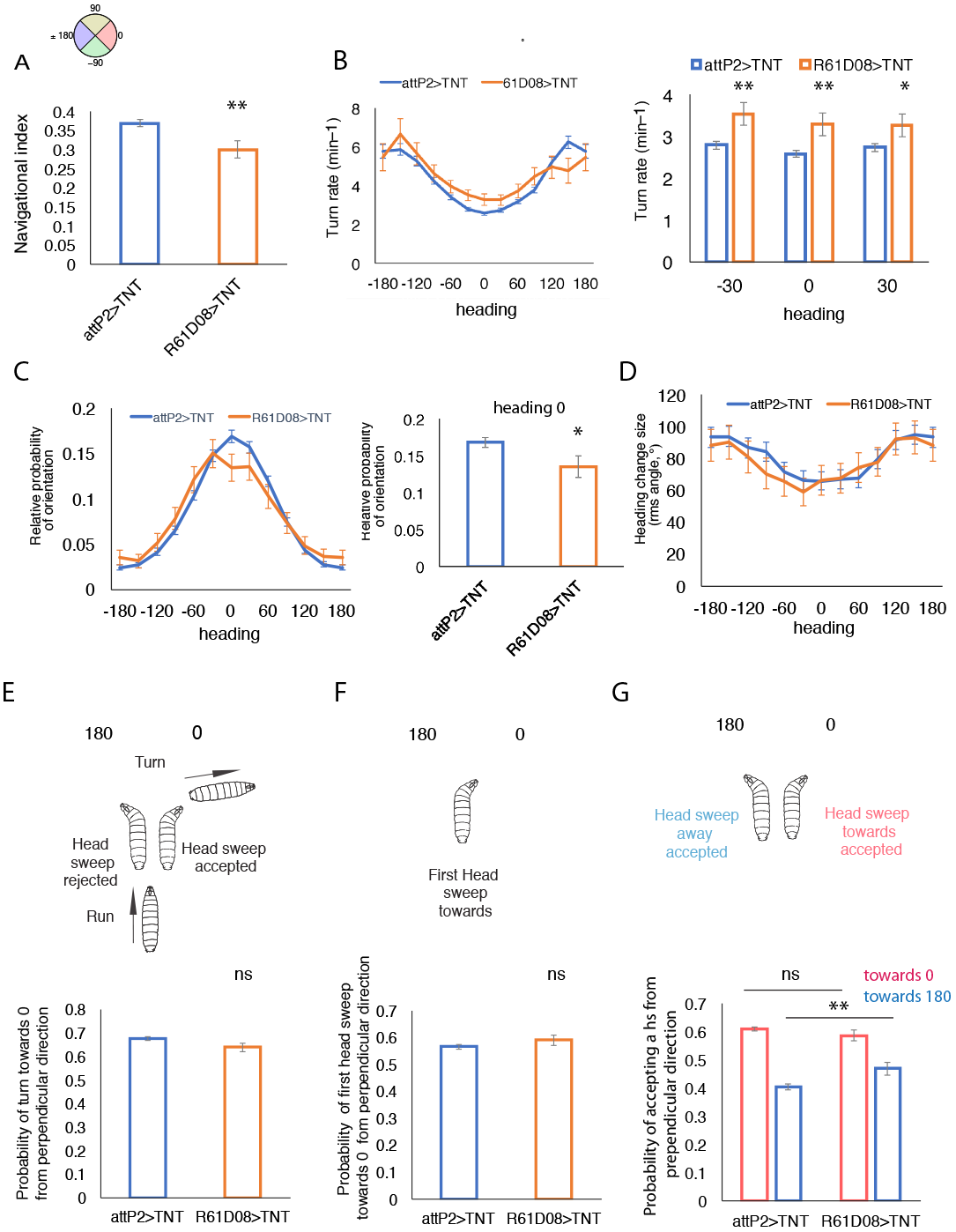
A. Comparison of the navigation index in chordotonal>TNT (R61D08>TNT) and attP2>TNT control larvae versus heading direction B. Comparison of turn rates in R61D08>TNT (chordotonal silenced) and attP2>TNT (control) larvae (left). Turn rate is higher in larvae with silenced chordotonal neurons when heading towards favorable directions (heading directions: −30, 0, 30) (right) C. Comparison of turn amplitudes in R61D08>TNT (chordotonal silenced) and attP2>TNT (control) larvae as a function of heading showed no significant difference D. Comparison of probability of orientation during runs in control attP2>TNT and R61D08>TNT larvae as a function of heading (left) and for heading towards 0 (right). E. Probability of turning towards 0 from perpendicular direction F. Probability of first headsweep towards 0 from perpendicular direction is not affected in larvae with silenced chordotonal sensory neurons G. Larvae with silenced chordotonal sensory neurons show a higher probability of accepting a headsweep towards 0 from perpendicular direction. mean and s.e.m, *: p<0.05, **p<0.01,**: p<0.001, p-values can be found in Supplementary tables 1-6.

Larvae with silenced chordotonal sensory neurons are defective in some of the strategies described above. They don’t decrease their turn rate when facing the lower speed end (heading towards direction from −30 to 30) as much as the control (Figure 3 B). Also, the turn amplitude as a function of heading is not significantly different compared to the control (Supplementary table 4, Figure 3 D). However, these larvae have lower probability of orientation towards the favorable direction compared to the control (heading towards −30 0 30) (Figure 3 C). The ability of larvae to bias the direction of their first headsweep, as well as the direction of the turn overall, is also not affected by silencing the chordotonal neurons (Figure 3 E-F, Supplementary table 3). However, the larvae with silenced chordotonal neurons have a higher probability of accepting a headsweep away from unfavorable direction (Figure 3 G) compared to the control. Overall, when chordotonal sensory neurons are silenced, anemotaxis is not completely impaired. We therefore tested another type of sensory neurons that was shown to mediate light touch and that we found in a previous study was involved in sensing air-current the multidendritic class III neurons [23], to see whether these neurons also are required for anemotaxis. However, silencing of multidendritic classs III neurons resulted in normal navigational performance (Supplementary figure 2 A).

### Identifying the neural substrates of anemotaxis

In order to identify key neurons in the central nervous system involved in anemotaxis, we further performed a targeted screen of 205 sparse GAL4 and intersectional Split GAL4 lines. These 205 lines were selected in an anatomical prescreen for lines that labeled brain neurons or candidate neurons in the somatosensory circuitry. These were identified in a previous study for mapping neurons underlying sensorimotor responses to air-puff (of uniform air-speeds) [23] and by matching light microscopy and EM reconstruction images [22, 24]. We generated the intersectional lines based on this prescreen in order to obtain even sparser neuronal expression patterns, with lines labeling single neuron types in many of the lines.

Because generally larvae are more efficient in navigation in stronger air-speed gradients (Figure 1 D, Supplementary table 1), we chose the 5 to 2 m/s gradient to perform the screen.

The high-throughput assay allowed us to accumulate enough larval trajectories (runs and turns) to discriminate the effects of small differences in larval navigational strategies upon neuronal manipulation.

We identified 8 neuronal lines in which the silencing of neurons resulted in less efficient taxis (their navigational index in the x axis was lower than the control) and 1 line in which the silencing of neurons resulted in more efficient taxis (R45D11) (the navigational index in x axis is higher than the control) (Figure 4 A and 7 A). For these lines, the navigational index in y axis was not significantly different from the control (except for SS00886) (Figure 4 B and 7 B). The number of animals tested and experiments performed is shown in Supplementary table 7.

**Figure 4.**
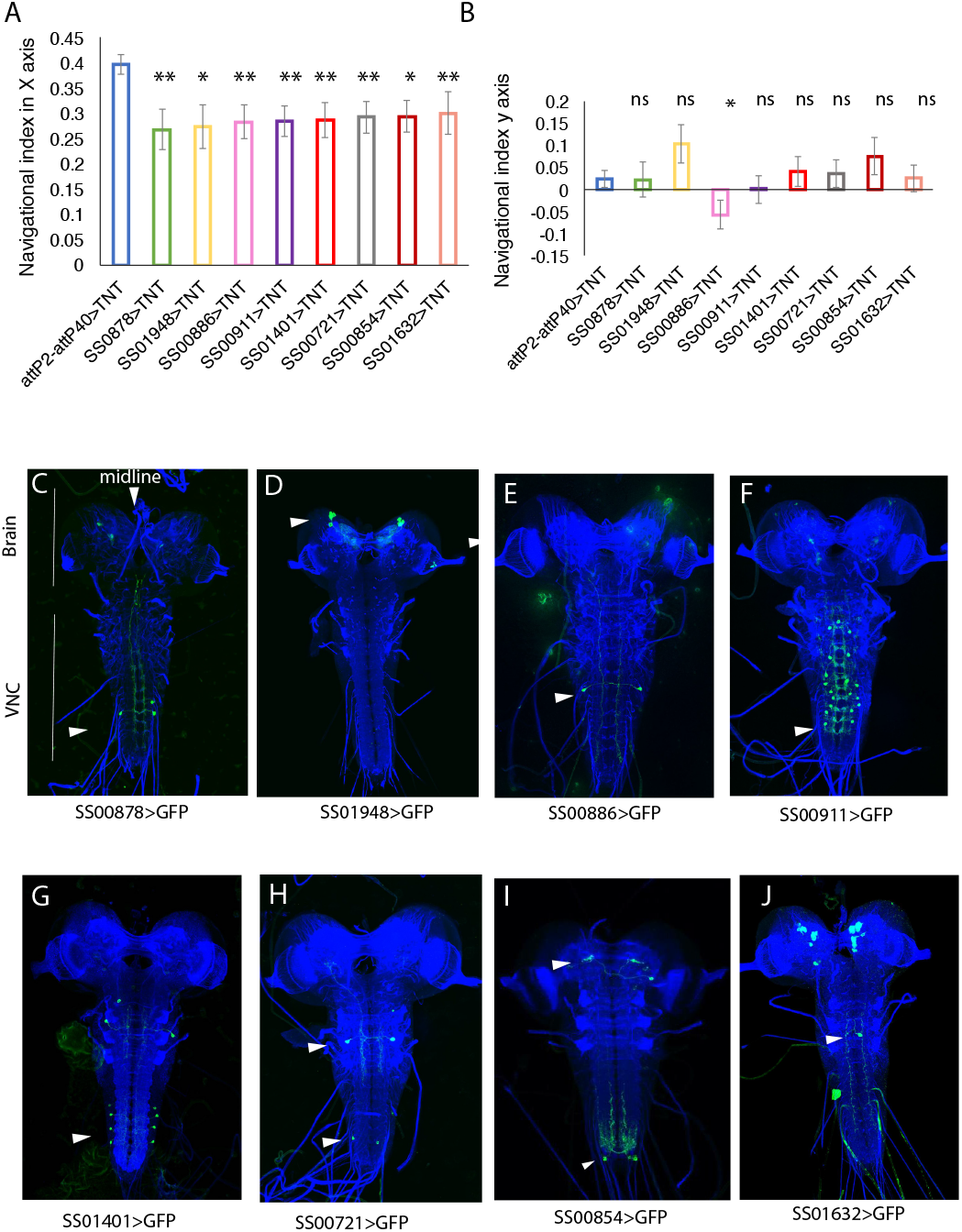
A. Navigational indices in X axis of 8 lines with less efficient anemotaxis compared to the control attP2-attP40>TNT. B. Navigational indices of 8 lines in Y axis with less efficient anemotaxis compared to the control attP2-attP40>TNT Mean and s.e.m are shown. *: p<0.05,**: p<0.01,***:<p<0.001 (p-values can be found in Supplementary table 2). C-J neuronal expression patterns in the 8 lines with less efficient anemotaxis. The name of the Split-GAL4 driver driving GFP expression is shown under each expression pattern image

We next examined what type of navigational strategy was affected in these lines that caused the poor or excellent navigational performances of these larvae: whether to turn, how much to turn or which way to turn. Amongst the 8 lines with less efficient taxis there were different groups of hits depending on what types of strategy was affected.

We first examined whether the 8 lines with poor navigational performance modulate the turn rate as a function of the sensory gradient less than the control. To test whether larvae with poor navigational performances decrease their turn rate less when heading in the favorable direction and increase their turn rate less when facing unfavorable directions, we compared the turning rates in the lines with lower navigational indices to the control when there were heading towards favorable direction (headings from −30 to 30, at −30, 0 and 30) and when they were heading towards unfavorable conditions (headings −150, 180, 150) (Figure 5 A-B).

**Figure 5.**
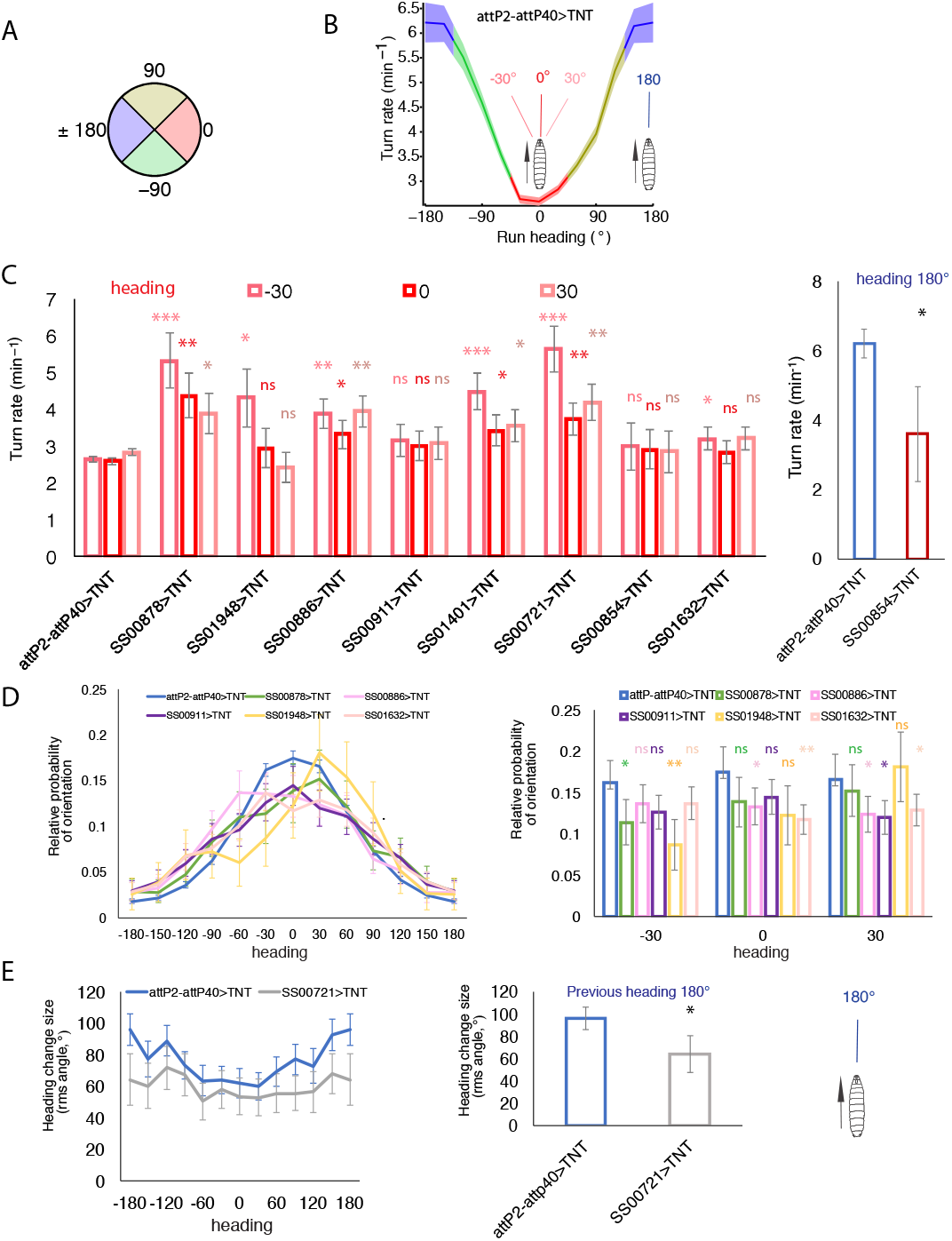
A. Compass indicating heading directions B. Turn rate versus heading for the attP2-attP40>TNT larvae. Schematics of larva head direction are shown on the plot C. Turn rates for larvae (lines as indicated) heading towards −30, 0 and 30 direction are shown on the left. Turn rate for 180 headings for SS00854>TNT larvae and the control on the right. D. Relative probability of headings during runs for attP2-attP40>TNT and five Split-GAL4 as indicated (left) and relative probability of headings during runs for directions heading towards −30, 0 and 30 (right) E. Turn size versus heading for the attp2-attP40>TNT and SS00721>TNT larvae (left). Comparison of the turn size when heading towards 180 in attP2-attP40>TNT and SS00721>TNT larvae (tight). values are means and s.e.m. *: p<0.05,**: p<0.01,***:<p<0.001 p-values can be found in Supplementary tables 5, 6 an 4 for D, E and F, respectively

In four (SS00721, SS00886, SS00878, SS01401) out of the eight lines the turn rate was increased in all the 3 favorable heading directions (-30, 0, 30), while the turn rate when facing unfavorable direction (180) was comparable to the control (Figure 5 C, Supplementary table 5). The lines SS001948 and SS01632 show significantly increased turn rate only when heading towards the −30 direction (Figure 5 C, Supplementary table 5).

The higher probability of turns when facing favorable directions would result in less runs towards favorable directions compared to the control. We measured this by computing the fraction of time that larvae spend crawling in different directions and found that the lines SS00886, SS00878, SS00911, SS001632 and SS01948 spent significantly less times heading towards lower wind speed end and showed a significantly lower probability of crawling orientation in at least one of the −30, 0, or 30 direction headings (Figure 5 D, Supplementary table 6). The lines SS00721 and SS01401 spent slightly, although not significantly, less time crawling in favorable directions (Supplementary table 6).

In one line, SS00854, we observed a decreased turn rate compared to the control when the larvae are facing unfavorable direction (Figure 5 C, right), which suggests that these larvae reorient less when facing unfavorable conditions and thus navigate less efficiently towards lower speed ends of the arena.

In addition to modulating the turn rate during navigation, larvae also modulate the amplitude of their turns so that they do smaller turns when heading towards favorable conditions and larger turns heading towards unfavorable conditions. We tested whether silencing in any of the 8 neuronal lines created a defect in this bias. We found that SS00721>TNT larvae modulate the turn amplitude less compared to the control (Figure 5 E, Supplementary table 4). When facing the unfavorable direction, the control larvae turn on average by 96 degrees, while SS00721>TNT larvae turn on average by 64 degrees (Figure 5 E).

We next examined how the directional decisions were affected in these lines with poor navigational performances. Seven out of the 8 lines have lower probabilities of turning towards the lower speed end from the orthogonal direction compared to the control (Figure 6 A, Supplementary table 3).

**Figure 6.**
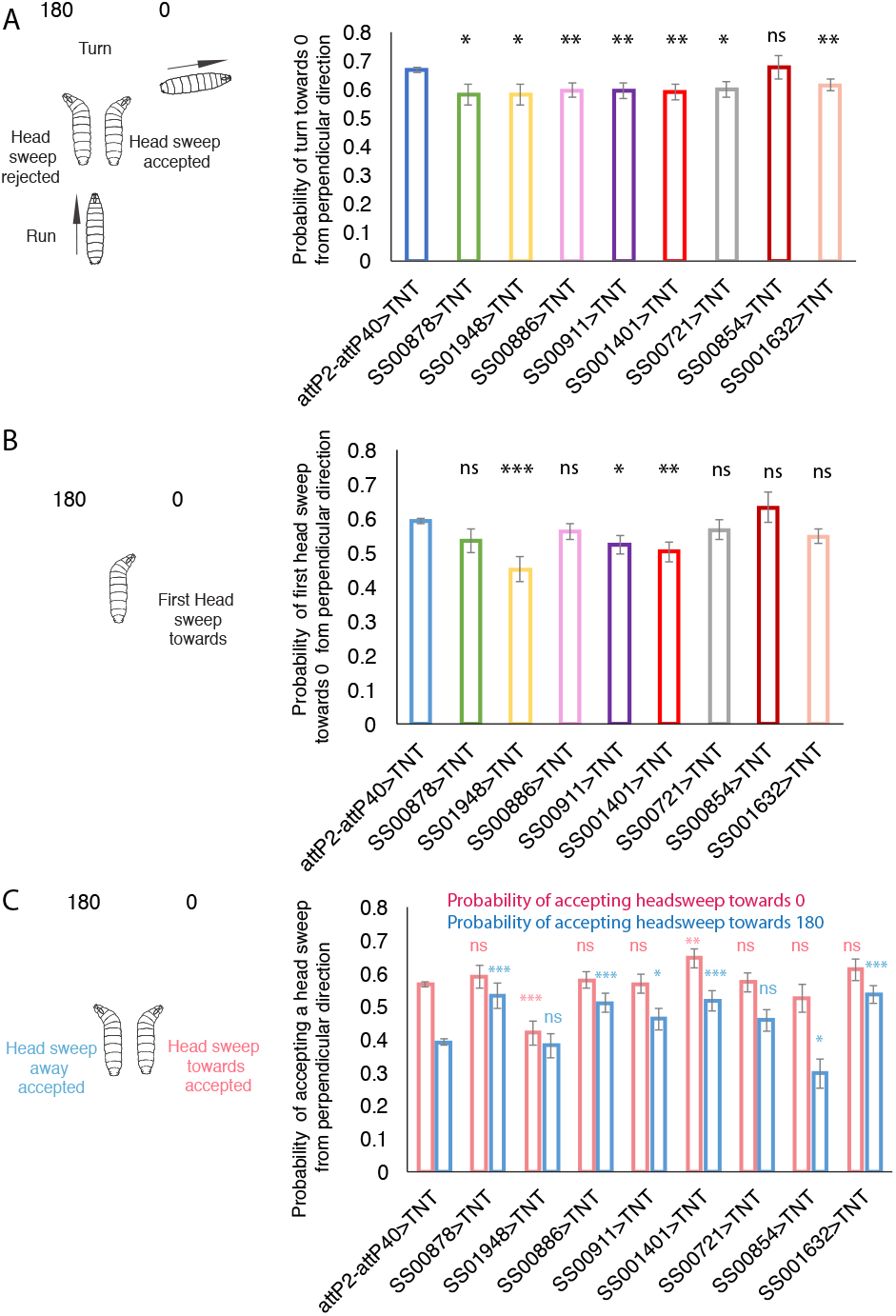
A. Probability of turns from perpendicular direction towards lower speed end (0) compared to the control attP2-attP40>TNT B. Probability of first head sweep from perpendicular direction towards lower speed end (0) compared to the control attP2-attP40>TNT C. Probability of starting a run during a head sweep from perpendicular direction compared to the control attP2-attP40>TNT. Mean and s.e.m derived from counting statistics. *: p<0.05,**: p<0.01,***:<p<0.001 (p-values can be found in Supplementary table 3)

As mentioned earlier third instar larvae show a bias of doing the first head-sweep towards the lower speed end during anemotaxis. The SS01948 line in which we silenced a pair of brain neurons, biases the probability of first head sweep toward the high-speed end instead (Figure 6 B, Supplementary table 3). In addition, these larvae show a lower probability of accepting head sweep towards 0 (favorable direction). None of the other 7 lines had a similar phenotype. Rather, the lines SS00878, SS00886, SS00911, SS01401 and SS01632 had the probability of accepting the head sweep and initiating a run in the 180 direction (unfavorable direction) significantly increased compared to the control (Figure 6 C, Supplementary table 3), suggesting the neurons in these lines normally suppress the initiation of runs from head sweeps in the unfavorable directions.

Out of 8 central nervous system (CNS) neuronal lines required for normal anemotaxis, 5 drive in single neuron types. The SS00721 labels 3 cells types stochastically: two pairs of thoracic neuron and a pair of abdominal neurons. One type of thoracic neuron was present in a majority of imaged larvae and is therefore very likely responsible for the phenotype observed. However, we cannot exclude that 2 other neuron type contribute to the observed phenotype.

Out of the single neuronal lines, we were able to identify the neurons of three by comparing light microscopy images to the EM reconstruction images (from previous studies, [22, 24]. Two neuronal lines drive in neurons downstream of nociceptive neurons, one ascending neuron A09o (SS00878) and one local abdominal neuron A10a (SS00911) that also receives input from basin-2 and basin-4 neurons [24]. The basin-2 and Basin-4 neurons where previously shown to be involved in avoidance response to mechanosensory and nociceptive responses [22, 24] and required for bending (head-cast or head sweep) in response to air-puff [22]. One line drive in neurons downstream of the chordotonal sensory neurons, SS01401, a local abdominal interneuron, drunken-1 [22]. Additional neuron types are also very likely downstream of somatosensory neurons: a thoracic descending neurons (SS001632), a thoracic descending neuron from the SS00721 line and an ascending neuron with a cell body in the abdominal a2 segments (SS00886), as their dendritic projections overlap with the axonal projections in the somatosensory domain (Supplementary figure 2 B-E) [25-27]. Two of the hit lines drive in two distinct brain neuron types: SS01948 and SS00854. In addition to the brain neurons, the SS00854 line also labels an ascending neuron with the cell body in the terminal abdominal segment (Figure 4 C-J). The SS01948 line drives gene expression in a cluster of dopaminergic neurons (DAN) that synapse onto the medial lobe of the mushroom body: DAN-i1, -j1, -k1 and -l1 [28]. These neurons were shown to mediate appetitive learning [29].

### Neurons that inhibit anemotaxis

We identified one line (R45D11) that exhibits more efficient taxis when driving tetanus-toxin (Figure 7 A).

**Figure 7.**
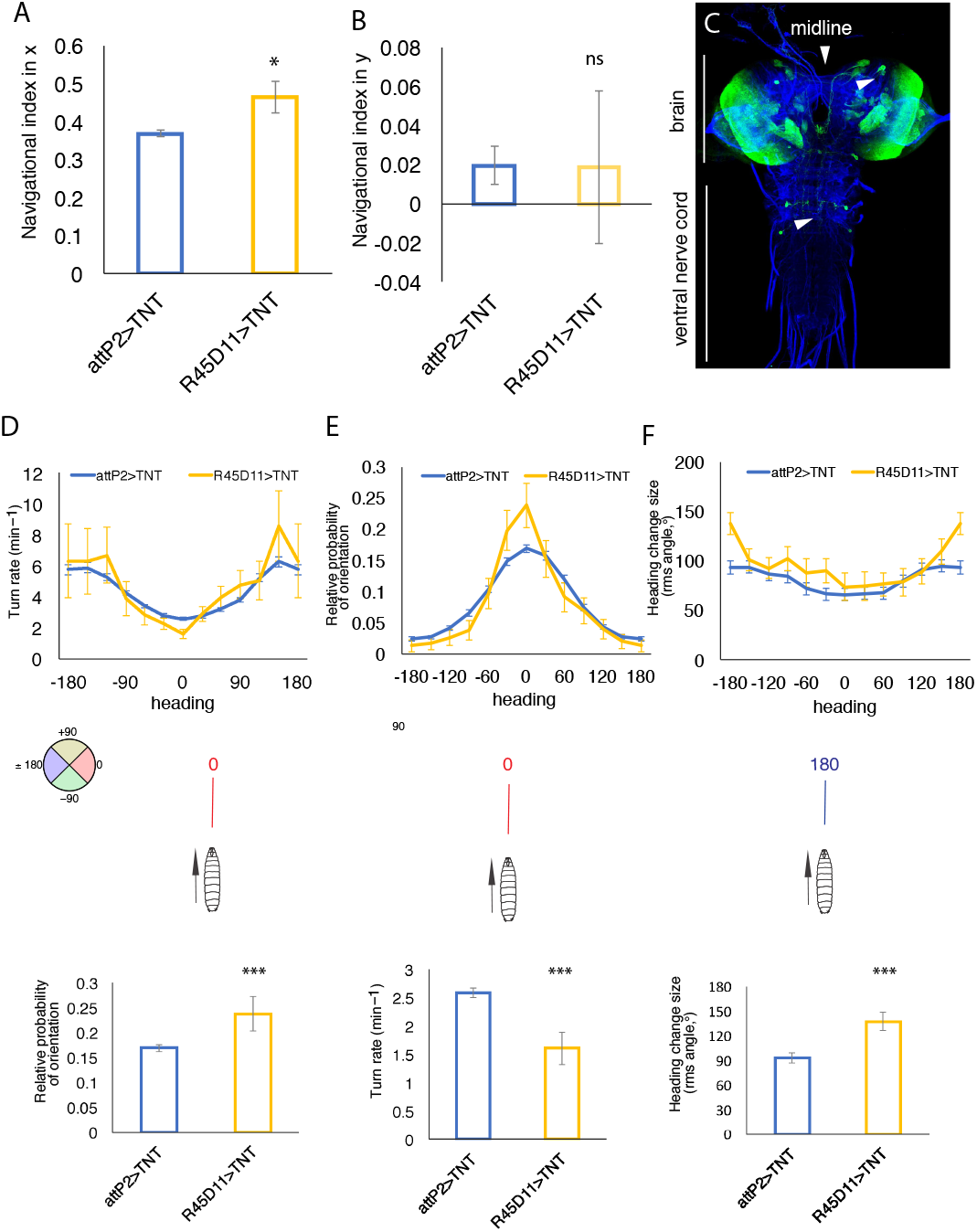
A. Navigational index for R45D11>TNT larvae with more efficient taxis in X axis compared to the control: attP2>TNT B. Navigational index for R45D11>TNT larvae with more efficient taxis in Y axis compared to the control: attP2>TNT C. Neuronal expression pattern for the R45D11 line D. Turn rate versus heading in all heading directions in R45D11>TNT and attP2>TNT larvae (top). Comparison of turn rates in the 0 heading direction in R45D11>TNT larvae and attP2>TNT (bottom) E. Relative probability of headings during runs in all direction (top) in attP2>TNT and R45D11>TNT larvae. Comparison of relative probability of headings during runs in R45D11>TNT larvae and attP2>TNT larvae in the 0 direction (bottom) F. Turn size versus heading in R45D11>TNT and attP2>TNT larvae (top) Comparison of turn size in the 0 heading direction in R45D11>TNT larvae and attP2>TNT (bottom) mean and s.e.m, p<0.05,**p<0.01,**: p<0.001, p-values can be found in Supplementary tables 2, 4-6

R45D11 is a sparse line that drives in a single pair of thoracic neurons and has sparse neuronal expression in the brain (Figure 7 C). Here we describe the observed phenotype. Overall the R45D11 larvae spend more time in runs towards the lower speed end and they are better in several navigational strategies compared to the control (Figure 7, Supplementary figure 3). For example, they increase the amplitude of turns significantly more when facing unfavorable conditions than the control, 138 compared to 93.5 degrees (Figure 7 F, Supplementary table 4), and they decrease their turn rate significantly more when facing favorable condition (Figure 7 E, Supplementary table 5), 1.6 compared to 2.6 turns per minute. They extend their runs significantly more when heading towards favorable conditions (Figure 7 D).

The directional decisions were less affected upon silencing (Supplementary figure 3 A-C). Only the probabilities of turning towards the favorable direction and accepting head sweep and initiating runs when facing favorable conditions is slightly higher (but not significantly) in R45D11 compared to the control, while the probability of accepting a head sweep towards or away from favorable conditions is unaffected (Supplementary figure 3).

Overall the R45D11>TNT larvae modulate their turns (turn rate and turn amplitude) and runs as a function of the air-current speeds more efficiently than the control. The directional decisions are unaffected by silencing and remain comparable to the control, which suggests that the navigational strategies involving decisions when and how much to turn are independently controlled from directional decisions (which way to turn).

## Discussion

We characterize for the first time the behavioral dynamics and the neural substrates underlying anemotaxis in *Drosophila* larvae navigating gradients of wind speed. We first determined that larvae move down the gradient, away from strong air-currents (winds). Using quantitative behavioral analysis, we then identified the behavioral strategies that larvae use to achieve this behavior and find that larvae use similar strategies previously described for larval navigation in other types of sensory gradients: periods of forwards crawls and turns that they modulate as a function of the sensory environment. We showed that chordotonal sensory neurons, previously known to mediate on-off responses to air current, also mediate anemotaxis.

By combining the analysis with manipulation of neuronal activity using tetanus-toxin (silencing) of very sparse neuronal population and single cell types in a targeted behavioral screen of 205 lines, we uncovered 8 neuronal lines driving expression in central neurons in the VNC and brain that are required for navigational decisions during anemotaxis. We described the phenotypes that result from silencing of these neurons using specific GAL4 drivers and Split GAL4 lines during anemotaxis and found that some strategies were affected in several hit lines and as a result of the manipulation of the activity of different neuron types (Supplementary table 8).

During navigation larvae increase their turning rate when facing unfavorable conditions and extend runs when facing favorable conditions.

Interestingly we find that some lines with lower navigational indices do not show a decrease in turning rate in unfavorable conditions but rather an increase of turning rate when heading towards favorable conditions compared to the control. This suggests that the increase of turning rate in unfavorable conditions and decrease in turning rate in favorable conditions are independently controlled. And this further suggests that larvae not only increase turn rate in response to unfavorable air current conditions, but also actively decrease turn rate in response to favorable air current conditions.

During reorientation events (turns) larvae do one or more head sweeps during which they perform temporal comparison of sensory information based on which they select a direction for the next run.

When examining the probability of accepting a head sweep when facing towards or away the favorable direction, we find that in lines that showed a phenotype in this strategy, generally the probability of accepting the head sweep and extending a run when facing the favorable direction (weak wind) were not lower in the lines we investigated (except in one line), but the probability of accepting a head sweep when facing the unfavorable direction (strong wind) was increased in 6 out of 8 lines, suggesting that these larvae, are, at least in part, less efficient in anemotaxis because they reject less head sweeps that don’t improve their condition. This could be due either to defects in sensory processing (of air-current speeds/intensity) or the inability to make appropriate directional decisions. Again, this points to independent control of acceptance of improving conditions and rejection of deteriorating conditions.

Larval somatosensory neurons cover larval body wall from head to tail [30, 31]. This is different from most other sensory neurons that mediate taxis behavior in other sensory gradients which are primarily located in the head [1, 4, 5, 9, 13, 14]. This might cause some difference in how comparisons are performed during head sweeps. Overall, we found that larvae use similar strategies during anemotaxis as they use during navigation in other types of sensory gradients. Previous studies showed that the first head sweep was unbiased in some other types of navigation, i.e in ethyl-acetate and C0_2_ gradients [5]. We find that in anemotaxis 59 % head sweep in attP2-attp40 control larvae and 56% of in attP2 control larvae are towards the favorable direction, when they are positioned perpendicularly to the gradient. This suggest that during anemotaxis larvae could be making spatial comparisons as a result of greater distance between sensory neurons along the body wall on the left and right side compared to the sensory neurons in the head. However, we cannot exclude that this difference is due to the difference stage of larvae used in previous studies (second instar) and current study (third instar). Furthermore, silencing of one type of somatosensory neurons, the chordotonal neurons, does not affect the probability of a favorable first head sweep.

In general, silencing chordotonals did not abolish anemotaxis completely and only affected the behavioral strategies moderately, which suggests that other types of somatosensory neurons also mediate anemotaxis. Since multidendritic class III don’t seem to be required for anemotaxis, the candidate sensory neuron types on the body wall are the external sensory neurons which by their morphology are well poised to mediate air-current related behaviors [30, 31].

We also identified neurons whose silencing resulted in unique phenotypes.

One line showed a decrease in turn rate when heading towards the high wind speed direction (SS00854) that was sufficient to result in lower navigational performance.

A strategy that larvae use during navigation in sensory gradients is to modulate the turn amplitude as a function of the gradient: they make smaller turns when facing the favorable direction and larger turns when facing unfavorable directions. Silencing of the neurons in the SS00721 neuronal lines affected not only the turn rate, but also the turn amplitude. These larvae perform turns of lower amplitudes when they have been heading towards stronger winds compared to the control, which would contribute to their poor navigational performance, as their reorientation away from strong winds will be less efficient.

That are two main categories of navigational decisions, those that pertain to the modulations of turns and runs and those that pertain to the choice of reorientation direction. We found one neuron type A10a (SS00911 line) the silencing of which didn’t affect the turn rate and turn amplitude, but the bias in the direction of turns in these larvae was significantly different from the control, which suggests that the two different types of decisions: when to turn and which way to turn, are independently controlled.

We also identified one neuronal line (R45D11) in which silencing of neuronal activity resulted in more efficient taxis. This is mainly achieved through a lower turning rate when heading towards lower speed direction and increased amplitude of turns when facing higher speed direction which results in more time spent crawling forward towards favorable conditions and more efficient reorientations respectively. Interestingly silencing these neurons had no effect on decisions affecting turn direction, further supporting the idea that these two types of strategies could be controlled separately. An increase in taxis efficiency upon silencing suggest that the neurons labeled by R45D11 might modulate the activities of neurons that normally promote navigational decisions. This might be useful in order to attend to other environmental conditions, like for example during foraging for food that is sensed through the presence of appetitive odors

Overall our findings support a model of navigation decision-making where different types of navigational decisions are controlled independently and are implemented by different modules in the nervous system.

In natural environments, taxis is rarely performed in the presence of stimuli from a single modality but rather sensory information from multiple modalities are combined and often larvae need to navigate in conflicting sensory gradients. In a previous study [8] we proposed that sensory information from multiple modalities are combined early in the navigational circuitry before the actual decision (to turn) point. In such an organization of the navigational circuitry the circuits that underlie navigational decisions (when to turn which way to turn and how much to turn) are likely to be shared between the navigational circuitries underlying different type of taxes. The idea of an overlapping circuitry of the different taxis behavior (navigations in different types of sensory gradients) is further supported by the findings that all the different taxis described so far (including anemotaxis described in the present study) use similar strategies to navigate towards more favorable environments. In line with this hypothesis is the recent finding that a group of neurons in the SEZ (subesophaegial zone) modulates the probability of transitioning between elementary behavior routines (runs and turns) based on information from multiple sensory modalities [16]. Identifying candidate elements of the navigational circuitry in higher order centers, i.e the brain (for example the SS00854 brain neurons and the mushroom body DAN neurons that we show here are involved in anemotaxis) that mediate navigational decisions in a gradient of air-speeds can therefore help elucidate the circuit mechanisms underlying not only anemotaxis, but navigational decision-making of Drosophila larvae in sensory gradients in general. For example, our findings suggest that the reward mediating dopaminergic neurons DAN-i1, -j1, -k1 and –l1 are implicated in navigational decision-making. These neurons synapse onto the medial lobe of the mushroom body, an integrative structure of the insect brain and are therefore very likely to be involved in decisionmaking in other modalities, not only during anemotaxis. In the adult *Drosophila,* DAN neurons show context and state movement related responses [32], while midbrain dopaminergic neurons in vertebrates were shown to be involved in movement initiation [33]. Thus, these dopaminergic neurons could be modulating the activity of higher-order centers in the mushroom body depending on the multisensory context or animal’s behavioral states [34] and thus could contribute to the sensorimotor decision-making (to turn or not to turn and which way to turn) during navigation in different modalities

Studying the neural basis of navigation in a genetically modifiable Drosophila larva has many advantages as neuronal activity can be manipulated, the connections between neurons determined by EM reconstruction [22, 24, 28, 35-40] and patterns of neuronal activity correlated with behaviors. The identified sparse and single neuron type lines in this study are excellent starting point for further studying the circuit mechanism underlying navigational decision-making by combining quantitative behavioral analysis with optogenetics and the monitoring of neuronal activity in a behaving animal [41].

## Methods details

### Drosophila Stocks

We used GAL4 from the Rubin collection available from Bloomington stock center each of which is associated with an image of the neuronal expression pattern shown at http://flweb.janelia.org/cgi-bin/flew. cgi. We used GAL4 line in behavioral experiments and generate intersectional lines (Split lines). In addition we used the insertion site stocks, w;attP2 and w;attP2;attP40 [42, 43], 19-12-GAL4, [44]. We used the progeny larvae from the insertion site stocks, w;;attp2, and w;attP2;attP40 crossed to the appropriate effector (UAS-TNT-e (II)) for characterizing the w;; attP2 and w;attP2;attP40 were selected because they have the same genetic background as the GAL4 and SplitGAL4 tested in the screen. We used the following effector stocks: UAS-TNT-e [45] and pJFRC12-10XUAS-IVSmyr::GFP (Bloomington stock number: 32197).

### Larva dissection and immunocytochemistry

To analyze the expression pattern of the GAL4 and SplitGAl4 lines, we crossed the lines to pJFRC12-10XUAS-IVSmyr::GFP (Bloomington stock number: 32197; [23]). The progeny larvae were placed in a phosphate buffered saline (PBS; pH 7.4) and fixed with 4.0% paraformaldehyde for 1-2 hr at room temperature, and then rinsed several times in PBS with 1% Triton X-100 (PBS-TX). Tissues when then mounted on poly-L-lysine(Sigma-Aldrich) coated coverslips and then transferred to a coverslip staining JAR (Electron Microscopy Sciences) with blocking solution, 3% normal donkey serum in PBS-TX for 1 hr. Primary antibodies were used at a concentration of 1:1000 for rabbit anti-GFP (Invitrogen) and 1:50 for mouse antineuroglian (Developmental Studies Hybridoma Bank) and 1:50 for anti-N-cadherin (Developmental Studies Hybridoma Bank) and incubated for 2 days at 4°C. Tissues when rinsed multiple times in PBS-TX and then incubated for 2 days. with secondary antibodies: anti-mouse IgG Alexa Fluor 568 Donkey (diluted 1:500; Invitrogen), Alexa Fluor 647 Donkey anti-rat IgG (1:500, Jackson ImmunoResearch) and fluorescein FITC conjugated Donkey anti-rabbit (diluted 1:500; Jackson ImmunoResearch). After incubation, the tissue was rinsed for several hours in PBSTX, and dehydrated through a graded ethanol series, cleared in xylene and mounted in DPX (Sigma) Images were obtained with 40x oil immersion objective (NA 1.3) on a Zeiss 510 Confocal microscope. Images of each nervous system were assembled from a 2xarray of tiled stacks, with each stack scanned as an 8 bit image with a resolution of 512×512 and a Z-step of 2 μm. Images were processed using Fiji (http://fiji.sc/) and ImageJ (https://imagej.nih.gov/ij/).

### Behavioral apparatus

The apparatus was described previously [21, 22]. Briefly, the apparatus comprises a video camera (DALSA Falcon 4M30 camera) for monitoring larvae, a ring light illuminator (Cree C503B-RCS-CW0Z0AA1 at 624 nm in the red), a computer (see [21]for details); available upon request are the bill of materials, schematic diagrams and PCB CAM files for the assembly of the apparatus) and a hardware modules for controlling air-puff, controlled through multi worm tracker (MWT) software (http://sourceforge.net/projects/mwt) [46], as described in [21]. Air-puff is delivered as described previously [21]. Briefly it is applied to a 25625 cm2 arena at a pressure of 1.1 MPa through a 3D-printed flare nozzle placed above the arena (with a 16 cm 6 0.17 cm opening) connected through a tubing system to plant supplied compressed air (0.5 MPa converted to a maximum of 1.4 MPa using a Maxpro Technologies DLA 5-1 air amplifier, standard quality for medical air with dewpoint of 210uC at 90 psig; relative humidity at 25uC and 32uC, ca. 1.2% and 0.9%, respectively). The strength of the airflow is controlled through a regulator downstream from the air amplifier and turned on and off with a solenoid valve (Parker Skinner 71215SN2GN00). The gradient is achieved by adjusting the inclination of the nozzle delivering the air-current to the arena. The gradient of airflow speeds is parallel to the direction of air-flow and decreases with the distance from the nozzle (source of wind) . The nozzle is fixed with a system of screws to prevent any movement during and in between experiments. Air-flow rates were measured before each round of experiments at 9 different equidistant positions in the arena with a hotwire anemometer to ensure that the speed was 5 m/s at one end and on the opposite end 2 m/s for the 5-2 m/s gradient and 3 m/s and 1 m/s respectively, for the 3-1 m/s gradient (Extech Model 407119A and Accusense model UAS1000 by DegreeC). The air-current relay is triggered through TTL pulses delivered by a Measurement Computing PCI-CTR05 5-channel, counter/timer board at the direction of the MWT. The onset and durations of the stimulus is also controlled through the MWT.

### Behavioral Experiments

Embryos were collected for 8–16 hours at 25°C with 65% humidity. Larvae were raised at 25°C with normal cornmeal food. Foraging 3^rd^ instar larvae were used (larvae reared 72-84 hours or for 3 days at 25°C).

Before experiments, larvae were separated from food using 10% sucrose, scooped with a paint brush into a sieve and washed with water (as described previously). This is because sucrose is denser than water, and larvae quickly float up in sucrose making scooping them out from food a lot faster and easier. This method is especially useful for experiments with large number of animals. We have controlled for the effect and have seen no difference in the behavior between larvae scooped with sucrose and larvae scooped directly from the food plate with a pair of forceps.

The larvae were dried and spread on the agar starting from the center of the arena. The substrate for behavioral experiments was a 3% Bacto agar gel in a 25625 cm2 square plastic dishes. Larvae were washed with water at room temperature, the dishes were kept at room temperature and the temperature on the rig inside the enclosure was set to 25°C.

The humidity in the room is monitored and held at 58%, with humidifiers (Humidifirst Mist Pac-5 Ultrasonic Humidifier).

We tested approximately 20–30 larvae at once in the behavioral assays. For each genotype, we did at least 3 repetitions (see Supplementary table 7 for N of animals and experiments) with at least 30 larvae total analyzed. The temperature of the entire rig was kept at 25 °C. In the assay, the larvae were put in the center of the plate in a line perpendicular to the gradient axis immediately prior the stimulus delivery. The air-puff was delivered continuously from the beginning of the experiment (time 2s after start of recording) and then for 10 minutes.

The N of animals and experiments is given in Supplementary table 7.

## BEHAVIORAL ANALYSIS

### Larva tracking

Larvae were tracked in real-time using the MWT software [46]. We rejected objects that were tracked for less than 5 seconds or moved less than one body length of the larva. For each larva MWT returns a contour, spine and center of mass as a function of time. From the MWT tracking data we computed the key parameters of larval motion, using the MAGAT analyzer software package (https://github.com/samuellab/MAGATAnalyzer) that we adapted to the MWT format [47]. Further analysis was carried out using custom MATLAB scripts described previously [47] software to identify behaviors, especially runs, turns, and head sweeps.

To calculate statistics involving center-of-mass movement along larval trajectories (for example, distributions of instantaneous heading and speed in Figures 2 and Supplementary Figure 1 and navigational indices in Figures 1, 3 and 6 and Supplementary figure 1) we needed to estimate the number of independent observations of quantities of center-of-mass movement along each larval trajectory. To do this, we calculated the autocorrelation function of the direction of motion,

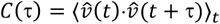

and extracted the time constant, T, of its component of exponential decay,

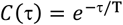

This correlation time constant was typically ∽20 s. To calculate the s.e.m. of center-of-mass motion statistics, we estimated the number of independent observations as the total observation time for each measurement divided by twice the correlation time constant. For more details see [47].

### Screen design

We screened a total of 205 Drosophila lines: 37 GAL4 lines from the Rubin GAL4 collection ([42, 48] and 168 Split GAL4lines made based on the GAL4 lines in the Rubin collection. We silenced small subsets of neurons and individual neurons in these lines using tetanus toxin. We selected these lines from the entire collection for sparse expression in the brain and ventral nerve cord of the larval CNS as well as expression in the sensory neurons (images of the larval CNS are available at http://www.janelia.org/gal4-gen1). The intersectional Split lines were designed based on the overlapping expression patterns of GAL4 driver lines. In addition for sparseness, some lines were chosen to be screened based on results from a previous air-puff inactivation screen (without gradient) [23].

The N of detected navigationals events (N or runs, reorientations) is given in Supplementary table 7.

### Hit detection and statistical analysis

The hits were determined based on the overall navigational performance of each GAL4 and Split-GAL4 lines. Hits were considered the lines that were significantly different compared to their respective controls w;;attP2 for GAL4 lines and w;attP2;attP40 for Split GAL4 lines. We used the normal distribution (Z) test to test the hypothesis that the navigational indices are the same at (rejection at p < 0.05 (without multiple comparisons).

We then analyzed the navigational strategies and compared the silencing experiments (for the 9 hit lines) with control experiments. We used the normal distribution (Z) test to test the hypothesis that modulation of turn rate and sizes is higher or equal unfavorable conditions and lower or equal in favorable conditions in lines with poorer navigational performances (based on the navigational index) and is lower or equal in unfavorable conditions and higher or equal in favorable conditions for the R45D11 line with more efficient navigation. Rejection is at p < 0.05.

We used the normal distribution (Z) test the hypothesis that the probabilities of turning towards 0, of accepting a head sweep towards 0 and towards 180 and of the first head sweep towards 0 are the same as the control (rejection at p < 0.05).

## Acknowledgments

We thank Yuh Nung Jan (UC San Francisco) for fly stocks. We thank Fly core (especially Monti Mercer) at JRC for fly crosses, Rebecca Arruda and Tam Dang for help with the behavioral experiments and Casey Schneider-Mizell for helpful discussion regarding neuronal connectivity. We thank Rex Kerr for assistance with MWT. We thank the Janelia Visitor Project for M. Gershow’s visits

## Supplementary Figure and table captions

**Supplementary Figure 1. -related to Figure 2.**
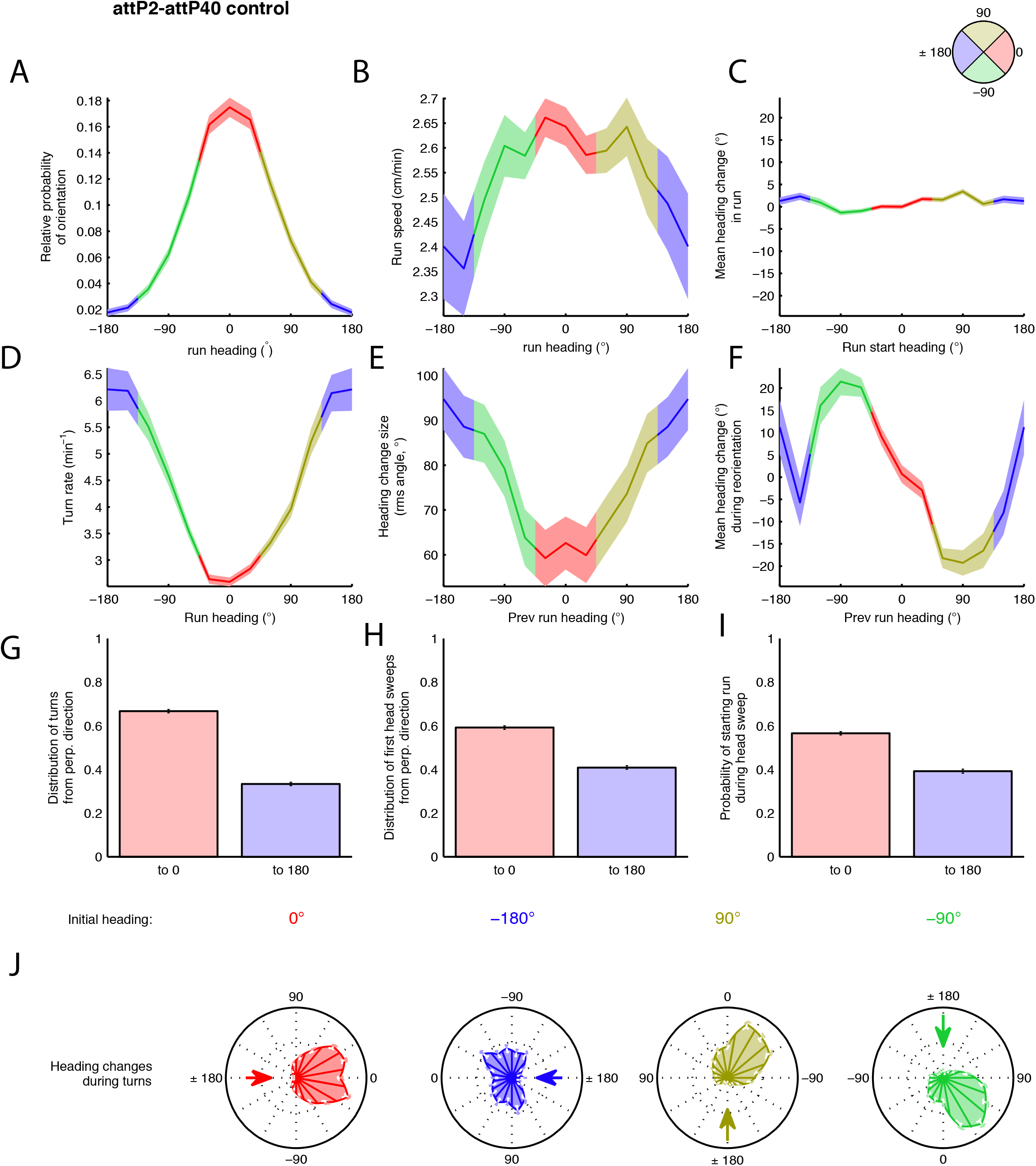
Navigational strategies in anemotaxis in control attP2-attP40>TNT A. Relative probability of headings during runs. Speed versus heading during runs B. Mean heading change in runs C. Turn rate versus heading D. Turn size versus heading E. Mean heading change during reorientation F. Distribution of turns from perpendicular direction G. Distribution of head sweeps from perpendicular direction H. Probability of starting a run during a headsweep I. Heading changes during runs sorted by initial heading J. Heading changes during runs sorted by initial heading. Values are mean and s.e.m

**Supplementary Figure 2. -related to Figures 3. and 4.**
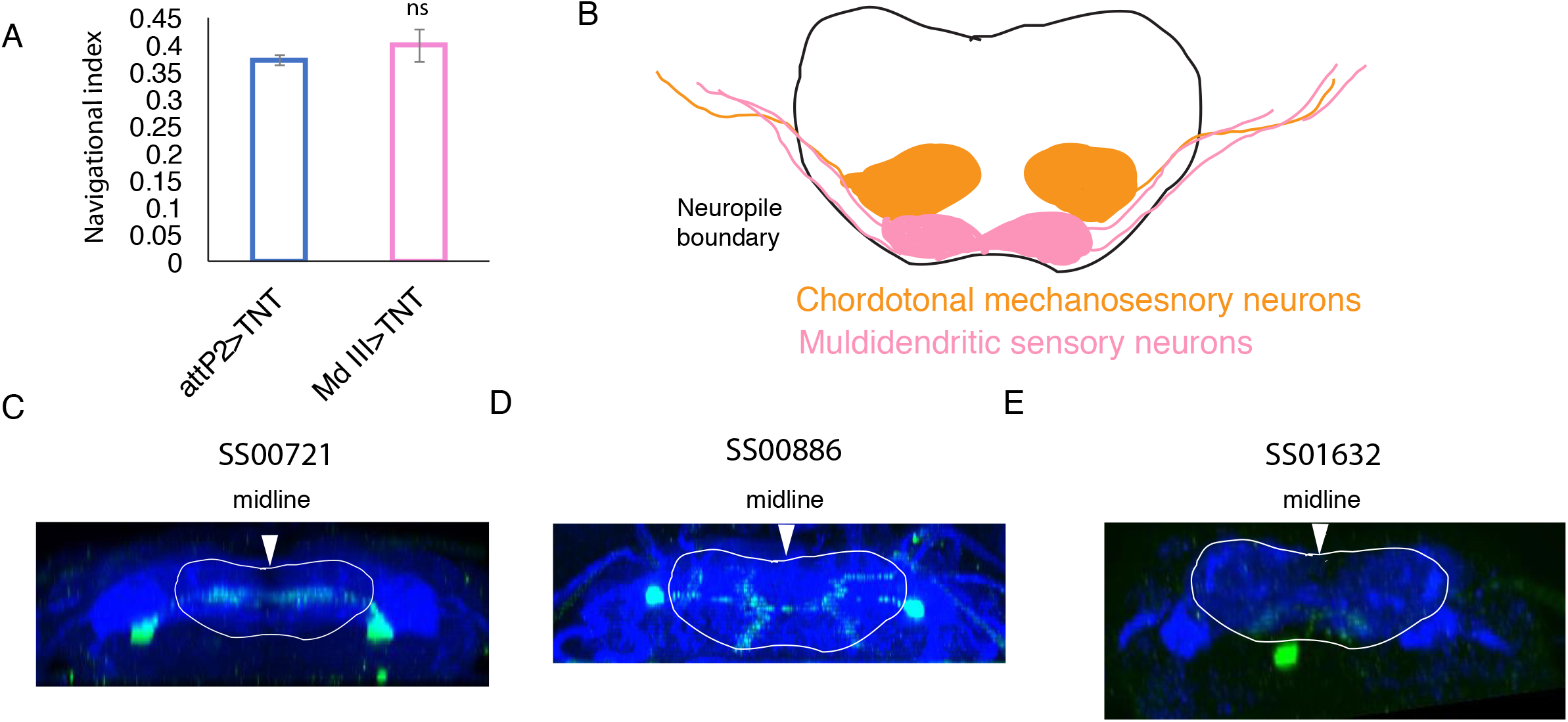
A. Navigational index for Md III>TNT larvae compared to control attP2>TNT larvae mean and s.e.m B. Schematics of projections in the VNC neuropil of mechanosensory and multidendritic sensory neurons A. SS00721>GFP, transverse projection of the thoracic region B. SS00886>GFP transverse projection in the abdominal region C. transversal section in the thoracic region SS01632>GFP. The dendrites of the neurons in A-C project to the somatosensory region of the Drosophila larva VNC neuropil that is rich in axonal projections from somatosensory neurons (projections for chordotonals sensory neurons-orange, multidendritic sensory neurons-pink) (1-3)

**Supplementary Figure 3. -related to Figure 7.**
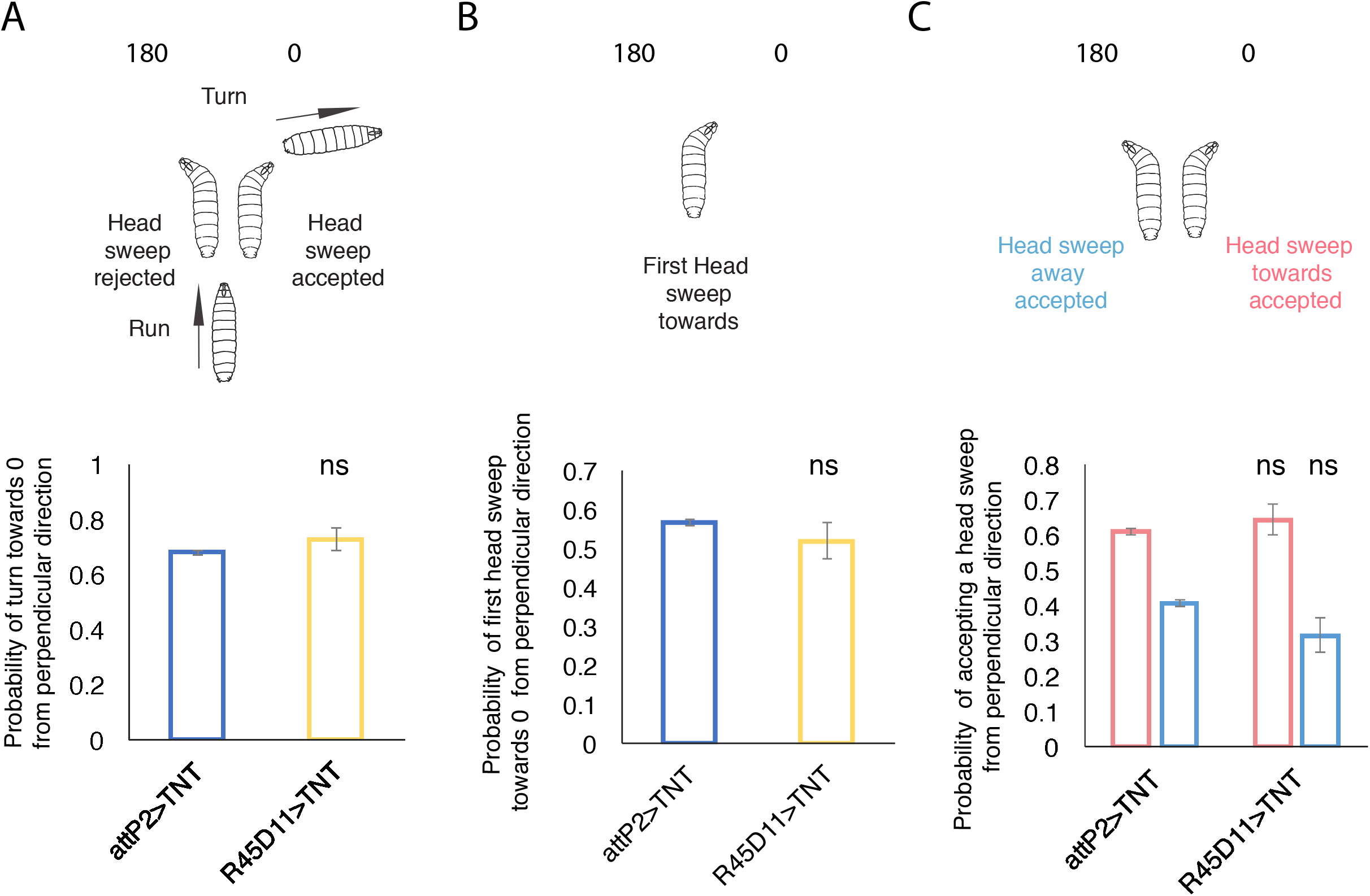
A. Probability of turns from perpendicular direction towards lower speed end (0) in R45D11>TNT compared to the control attP2>TNT B. Probability of first head sweeps from perpendicular direction towards lower speed end (0) in R45D11>TNT compared to the control attP2>TNT C. Probability of starting a run during a headsweep from perpendicular direction in R45D11>TNT compared to the control attP2>TNT. Mean and s.e.m are shown, p<0.05,**p<0.01,**: p<0.001, p-values can be found in Supplementary tables 3

**Supplementary table 1.** Navigational indices for sensory lines and control

**Supplementary table 2.** Navigational indices for central neuronal lines

**Supplementary table 3.** Directional decisions

**Supplementary table 4.** Turn size

**Supplementary table 5.** Turn rate

**Supplementary table 6.** Probability of orientation during runs

**Supplementary table 7.** Summary table for all experiments

**Supplementary table 8** Summary of phenotypes for hit GAL4 and Split-GAL4 lines. Low (wind speeds) represents any of the −30, 0, 30 direction, and high (wind speed) represents the 180 heading direction. < decreased, > increased, ns-non significant compared to the control.

## References

[1] Louis M, Huber T, Benton R, Sakmar TP, Vosshall LB. Bilateral olfactory sensory input enhances chemotaxis behavior. Nat Neurosci 2008;11:187–99. doi:10.1038/nn2031.

[2] Lockery SR. The computational worm: spatial orientation and its neuronal basis in C. elegans. Current Opinion in Neurobiology 2011;21:782–90. doi:10.1016/j.conb.2011.06.009.

[3] Lahiri S, Shen K, Klein M, Tang A, Kane E, Gershow M, et al. Two alternating motor programs drive navigation in Drosophila larva. PLoS ONE 2011;6:e23180. doi:10.1371/journal.pone.0023180.

[4] Kane EA, Gershow M, Afonso B, Larderet I, Klein M, Carter AR, et al. Sensorimotor structure of Drosophila larva phototaxis. Proc Natl Acad Sci USa 2013;110:E3868–77. doi:10.1073/pnas. 1215295110.

[5] Gershow M, Berck M, Mathew D, Luo L, Kane EA, Carlson JR, et al. Controlling airborne cues to study small animal navigation. Nat Methods 2012;9:290–6. doi:10.1038/nmeth.1853.

[6] Gomez-Marin A, Stephens GJ, Louis M. Active sampling and decision making in Drosophila chemotaxis. Nat Commun 2011;2:441. doi:10.1038/ncomms1455.

[7] Schulze A, Gomez-Marin A, Rajendran VG, Lott G, Musy M, Ahammad P, et al. Dynamical feature extraction at the sensory periphery guides chemotaxis. eLife 2015;4:1129. doi:10.7554/eLife.06694.

[8] Gepner R, Mihovilovic Skanata M, Bernat NM, Kaplow M, Gershow M. Computations underlying Drosophila phototaxis, odor-taxis, and multi-sensory integration. eLife 2015;4. doi:10.7554/eLife.06229.

[9] Luo L, Gershow M, Rosenzweig M, Kang K, Fang-Yen C, Garrity PA, et al. Navigational decision making in Drosophila thermotaxis. J Neurosci 2010;30:4261–72. doi:10.1523/JNEUROSCI.4090-09.2010.

[10] Wystrach A, Lagogiannis K, Webb B. Continuous lateral oscillations as a core mechanism for taxis in Drosophila larvae. eLife 2016;5:417. doi:10.7554/eLife.15504.

[11] Zhao W, Gong C, Ouyang Z, Wang P, Wang J, Zhou P, et al. Turns with multiple and single head cast mediate Drosophila larval light avoidance. PLoS ONE 2017;12:e0181193. doi:10.1371/journal.pone.0181193.

[12] Fishilevich E, Domingos AI, Asahina K, Naef F, Vosshall Jovanic et al., 07 JAN 2018 - preprint copy - BioRxiv LB, Louis M. Chemotaxis behavior mediated by single larval olfactory neurons in Drosophila. Curbio 2005;15:2086–96. doi:10.1016/j.cub.2005.11.016.

[13] Gomez-Marin A, Duistermars BJ, Frye MA, Louis M. Mechanisms of odor-tracking: multiple sensors for enhanced perception and behavior. Front Cell Neurosci 2010;4:6. doi:10.3389/fncel.2010.00006.

[14] Klein M, Afonso B, Vonner AJ, Hernandez-Nunez L, Berck M, Tabone CJ, et al. Sensory determinants of behavioral dynamics in Drosophila thermotaxis. Proc Natl Acad Sci USa 2015;112:E220–9. doi:10.1073/pnas.1416212112.

[15] Klein M, Krivov SV, Ferrer AJ, Luo L, Samuel AD, Karplus M. Exploratory search during directed navigation in C. elegans and Drosophila larva. eLife 2017;6:e30503–14. doi:10.7554/eLife.30503.

[16] Tastekin I, Riedl J, Schilling-Kurz V, Gomez-Marin A, Truman JW, Louis M. Role of the subesophageal zone in sensorimotor control of orientation in Drosophila larva. Curr Biol 2015;25:1448–60. doi:10.1016/j.cub.2015.04.016.

[17] Slater G, Levy P, Chan KLA, Larsen C. A central neural pathway controlling odor tracking in Drosophila. J Neurosci 2015;35:1831–48. doi:10.1523/JNEUROSCI.2331-14.2015.

[18] Spencer Johnston J. Genetic variation for anemotaxis (wind-directed movement) in laboratory and wild-caught populations Drosophilia. Behavior Genetics 1981;12.

[19] Kalmus H. Anemotaxis in Drosophila. Nature 1942; 150:405–1. doi:10.1038/150405a0.

[20] Flugge C. Geruchliche Raumorientierung von Drosophila melanogaster 1934.

[21] Ohyama T, Jovanic T, Denisov G, Dang TC, Hoffmann D, Kerr RA, et al. High-Throughput Analysis of Stimulus-Evoked Behaviors in Drosophila Larva Reveals Multiple Modality-Specific Escape Strategies. PLoS ONE 2013;8:e71706. doi:10.1371/journal.pone.0071706.

[22] Jovanic T, Schneider-Mizell CM, Shao M, Masson J-B, Denisov G, Fetter RD, et al. Competitive Disinhibition Mediates Behavioral Choice and Sequences in Drosophila. Cell 2016;167:858–870.e19. doi:10.1016/j.cell.2016.09.009.

[23] Jovanic T, Masson J-B, Truman JW, Zlatic M. Mapping neurons and brain regions underlying sensorimotor decisions and sequences in Drosophila. bioRxiv 2017:1–22. doi:10.1101/215236.

[24] Ohyama T, Schneider-Mizell CM, Fetter RD, Aleman JV, Franconville R, Rivera-Alba M, et al. A multilevel multimodal circuit enhances action selection in Drosophila. Nature 2015;520:633–9.

[25] Zlatic M, Li F, Strigini M, Grueber W, Bate M. Positional Cues in the Drosophila Nerve Cord: Semaphorins Pattern the Dorso-Ventral Axis. PLoS Biol 2009;7:e1000135. doi:10.1371/journal.pbio.1000135.

[26] Grueber WB, Ye B, Yang C-H, Younger S, Borden K, Jan LY, et al. Projections of Drosophila multidendritic neurons in the central nervous system: links with peripheral dendrite morphology. Development 2007;134:55–64. doi:10.1242/dev.02666.

[27] Merritt DJ, Whitington PM. Central projections of sensory neurons in the Drosophila embryo correlate with sensory modality, soma position, and proneural gene function. J Neurosci 1995;15:1755–67.

[28] Eichler K, Li F, Litwin-Kumar A, Park Y, Andrade I, Schneider-Mizell CM, et al. The complete connectome of a learning and memory centre in an insect brain. Nature 2017;548:175–82. doi:10.1038/nature23455.

[29] Rohwedder A, Wenz NL, Stehle B, Huser A, Yamagata N, Zlatic M, et al. Four Individually Identified Paired Dopamine Neurons Signal Reward in Larval Drosophila. Curr Biol 2016;26:661–9. doi:10.1016/j.cub.2016.01.012.

[30] Bodmer R, Jan Y-N. Morphological differentiation of the embryonic peripheral neurons in Drosophila. Rouxs Arch Dev Biol 1987;196:69–77. doi:10.1007/BF00402027.

[31] Merritt DJ, Whitington PM. Central projections of sensory neurons in the Drosophila embryo correlate with sensory modality, soma position, and proneural gene function. J Neurosci 1995;15:1755–67.

[32] Cohn R, Morantte I, Ruta V. Coordinated and Compartmentalized Neuromodulation Shapes Sensory Processing in Drosophila. Cell 2015;163:1742–55. doi:10.1016/j.cell.2015.11.019.

[33] da Silva JA, Tecuapetla F, Paixão V, Costa RM. Dopamine neuron activity before action initiation gates and invigorates future movements. Nature 2018;79:368. doi:10.1038/nature25457.

[34] Lewis LPC, Siju KP, Aso Y, Friedrich AB, Bulteel AJB, Rubin GM, et al. A Higher Brain Circuit for Immediate Integration of Conflicting Sensory Information in Drosophila. Curr Biol 2015;25:2203–14. doi:10.1016/j.cub.2015.07.015.

[35] Schneider-Mizell CM, Gerhard S, Longair M, Kazimiers T, Li F, Zwart MF, et al. Quantitative neuroanatomy for connectomics in Drosophila. eLife 2016;5:1133. doi:10.7554/eLife.12059.

[36] Fushiki A, Zwart MF, Kohsaka H, Fetter RD, Cardona A, Nose A, et al. A circuit mechanism for the propagation of waves of muscle contraction in Drosophila. eLife 2016;5:e13253. doi:10.7554/eLife.13253.

[37] Berck ME, Khandelwal A, Claus L, Hernandez-Nunez L, Si G, Tabone CJ, et al. The wiring diagram of a glomerular olfactory system. 2016. doi:10.1101/037721.

[38] Schlegel P, Texada MJ, Miroschnikow A, Schoofs A, Hückesfeld S, Peters M, et al. Synaptic transmission parallels neuromodulation in a central food-intake circuit. eLife 2016;5:462. doi:10.7554/eLife.16799.

[39] Zwart MF, Pulver SR, Truman JW, Fushiki A, Fetter RD, Cardona A, et al. Selective Inhibition Mediates the Sequential Recruitment of Motor Pools. Neuron 2016;91:615–28. doi:10.1016/j.neuron.2016.06.031.

[40] Larderet I, Fritsch PM, Gendre N, Neagu-Maier GL, Fetter RD, Schneider-Mizell CM, et al. Organization of the Drosophila larval visual circuit. eLife 2017;6:e28387–23. doi:10.7554/eLife.28387.

[41] Karagyozov D, Mihovilovic Skanata M, Lesar A, Gershow M. Recording neural activity in unrestrained animals with 3D tracking two photon microscopy. bioRxiv 2017:1–34. doi:10.1101/213942.

[42] Pfeiffer BD, Jenett A, Hammonds AS, Ngo T-TB, Misra S, Murphy C, et al. Tools for neuroanatomy and neurogenetics in Drosophila. Proc Natl Acad Sci USa 2008;105:9715–20. doi:10.1073/pnas.0803697105.

[43] Pfeiffer BD, Ngo T-TB, Hibbard KL, Murphy C, Jenett A, Truman JW, et al. Refinement of tools for targeted gene expression in Drosophila. Genetics 2010;186:735–55. doi:10.1534/genetics.110.119917.

[44] Zhang W, Yan Z, Jan LY, Jan Y-N. Sound response mediated by the TRP channels NOMPC, NANCHUNG, and Jovanic et al., 31. JAN 2018 INACTIVE in chordotonal organs of Drosophila larvae. Proc Natl Acad Sci USa 2013;110:13612–7. doi:10.1073/pnas.1312477110.

[45] Sweeney ST, Broadie K, Keane J, Niemann H, O’Kane CJ. Targeted expression of tetanus toxin light chain in Drosophila specifically eliminates synaptic transmission and causes behavioral defects. Neuron 1995;14:341–51.

[46] Swierczek NA, Giles AC, Rankin CH, Kerr RA. High-throughput behavioral analysis in C. elegans. Nat Methods 2011;8:592–8. doi:10.1038/nmeth.1625.

[47] Gershow M, Berck M, Mathew D, Luo L, Kane EA, Carlson JR, et al. Controlling airborne cues to study small animal navigation. Nat Methods 2012;9:290–6. doi:10.1038/nmeth.1853.

[48] Jenett A, Rubin GM, Ngo T-TB, Shepherd D, Murphy C, Dionne H, et al. A GAL4-Driver Line Resource for Drosophila Neurobiology. Cell Rep 2012;2:991–1001. doi:10.1016/j.celrep.2012.09.011.

## References

1. Zlatic M, Li F, Strigini M, Grueber W, Bate M. Positional Cues in the Drosophila Nerve Cord: Semaphorins Pattern the Dorso-Ventral Axis. Luo L, editor. PLoS Biol. Public Library of Science; 2009 Jun 23;7(6):e1000135.

2. Grueber WB, Ye B, Yang C-H, Younger S, Borden K, Jan LY, et al. Projections of Drosophila multidendritic neurons in the central nervous system: links with peripheral dendrite morphology. Development. The Company of Biologists Limited; 2007 Jan;134(1):55-64.

3. Merritt DJ, Whitington PM. Central projections of sensory neurons in the Drosophila embryo correlate with sensory modality, soma position, and proneural gene function. J Neurosci. Society for Neuroscience; 1995 Mar;15(3 Pt 1):1755-67.

